# Holocene plant diversity dynamics shows a distinct biogeographical pattern in temperate Europe

**DOI:** 10.1101/2020.08.19.257584

**Authors:** Jan Roleček, Vojtěch Abraham, Ondřej Vild, Helena Svitavská Svobodová, Eva Jamrichová, Zuzana Plesková, Petr Pokorný, Petr Kuneš

## Abstract

**Aims:** Reconstruction of the Holocene diversity changes in a biogeographically complex region. Description of major diversity patterns, testing their predictors, and their interpretation in the palaeoecological and biogeographical context. Testing the assumption that pollen record is informative with respect to plant diversity in our study area.

**Methods:** Fossil pollen extracted from 18 high-quality profiles was used as a proxy of past plant diversity. Pollen counts of tree taxa were corrected by pollen productivity, and pollen assemblages were resampled to 100 grains per sample and 150 grains per 500-years time window. SiZer analysis was used to test and visualize multi-scale diversity patterns. Linear modelling was used to identify the best predictors. SiZer maps and pollen composition were analysed using non-metric multidimensional scaling. K-means clustering and indicator species analysis were used to interpret ordination results.

**Results:** Mean Holocene plant diversity is significantly predicted by latitude, while its temporal pattern followed the biogeographical region and elevation. Major differences were found between the Mesic and Montane Hercynia (lower diversity, increasing only in the Late Holocene) and Pannonia, the Carpathians and Warm Hercynia (higher diversity, increasing from the Early or Middle Holocene on). Low diversity in the Middle and Late Holocene is associated with the prevalence of woody and acidophilic taxa. High diversity is associated with numerous grassland and minerotrophic wetland taxa, crops and weeds. Fossil-modern pollen diversity and modern pollen-plant diversity show significant positive relationships.

**Conclusions:** Plant diversity and its changes during the Holocene are geographically structured across temperate Europe. Main causes appear to be differences in past dynamics of the landscape openness and vegetation composition, driven mainly by changes in climate and human impact and their different timing. Fossil pollen, if appropriately treated, is a useful proxy of past plant diversity.

## Introduction

Diversity of life on the Earth has been changing over time, and biogeographers had paid attention to these changes long before the term “biodiversity” was coined (Blandin, 2015; Shmida & Wilson, 1985). Historical biogeography explicitly refers to the temporal aspect of biodiversity, and dynamic concept of species distributions, shaped e.g. by climate and migrations, was incorporated into the modern biogeographical thinking no later than in the seminal works of A. von Humboldt and A. de Candolle (Lomolino et al., 2004). Development of palaeoecology significantly improved our knowledge of the Quaternary dynamics of biodiversity and strengthened the vital links between biogeography and related disciplines such as ecology, evolutionary biology and palaeontology (Birks, 2019).

Holocene, the geological period covering the last 11,700 years, is a key time frame for understanding the present biodiversity (Birks, Felde & Seddon, 2016; Roberts, 2014). During the preceding 2.6 million years of the Quaternary, climatic fluctuations have driven global cycles of profound environmental changes: the glacial-interglacial cycles (Elias & Mock, 2013). In temperate Europe, these changes involved major shifts in biomes, with cold steppe, tundra and coniferous forest (taiga) being characteristic for the glacials and broad-leaved forests dominating the interglacials (Iversen 1958; Reille et al., 2000). The last such shift started by the end of the Last Glacial Maximum (around 20,000 cal BP) and was almost completed at the onset of the Middle Holocene (8200 cal BP). However, between 8000 and 7000 cal BP, Neolithic revolution engulfed temperate Europe, and humans became the key players in transformations of the environment (Kalis et al., 2003; Roberts, 2014). The constant changes had great effects on biotic assemblages (šizling et al., 2016) and the recent biodiversity change, connected to human activities (Pereira et al., 2012), may also be understood in this context (Birks, Felde & Sedddon, 2016).

Fossil pollen has been used as a proxy of past plant composition for more than 100 years (Edwards et al. 2017, Birks & Berglund 2018). However, concerns have been raised whether the pollen record is informative also concerning plant richness and diversity (Goring et al. 2013). Indeed, there are several sources of bias in the pollen-plant diversity relationship that might invalidate fossil pollen as a proxy. First, numerous plant species belonging to large taxonomic groups such as grasses, sedges or the Chenopodiaceae family, have morphologically similar pollen that cannot be distinguished by standard methods of pollen analysis. This allows coarser taxonomic resolution of pollen than that of plants. Second, pollen diversity estimates strongly depend on pollen sample size, and it is important to distinguish true absences from false absences due to rarity and limited pollen counts. Third, even in samples with equal pollen sums, detection probability of rare pollen types is influenced by abundant taxa and their skewed representation due to pollen production, dispersal, sedimentation and spatial arrangement of source plants (Odgaard, 1999).

Despite these complexities, results of recent palynological studies suggest that if adequately analysed, pollen richness shows good correspondence with plant richness (Felde et al., 2016; Meltsov et al., 2011; Reitalu et al., 2019). Also, a recent synthesis of fossil pollen data covering most of Europe (Giesecke et al., 2019) has shown apparent differences in the Holocene diversity levels and temporal trends at the continental scale. Existence of such coarse-scale differences is in agreement with common knowledge of pollen analysts about uneven levels of fossil pollen diversity across biomes (Birks, Felde, Bjune et al., 2016). However, there is a lack of evidence whether pollen allows detection also of finer-scale spatial patterns of diversity, e.g. within a single region. In other words, are differences in plant diversity among landscapes, which are of interest for biogeographers and ecologists, detectable in the fossil pollen record? A positive answer would open new opportunities for insight into the origin of the present plant diversity and for setting the temporal context for the current biodiversity change.

This study aims at the reconstruction of the Holocene diversity patterns in a single, biogeographically diverse region of temperate Europe. We describe major patterns, test their predictors and investigate how they can be interpreted using the available knowledge. Moreover, we test the assumption that pollen record is informative with respect to plant diversity in our study area, based on original pollen and plant diversity data.

## Materials and methods

### Study area

The study area covers approximately 110 thousand km^2^ and is situated in a part of temperate Europe that is complex in terms of topography and biogeography (Figure 1; Chytrý et al., 2017). It is also well studied from the palaeoecological point of view (Kuneš et al. 2009). The European Central Uplands form the western part of the study area. They consist of a ring of mountains, encircling relatively warm and dry basins and uplands. They are underlain predominantly by old crystalline bedrock, in the lowlands usually covered by younger calcareous sediments. Lower elevations generally have a long history of agricultural use (usually since the Neolithic), while higher elevation with acidic soils have been deforested mostly in the Middle Ages or remain forested up to now (Kozáková et al., 2015; Pokorný et al., 2015).

**Figure 1.**
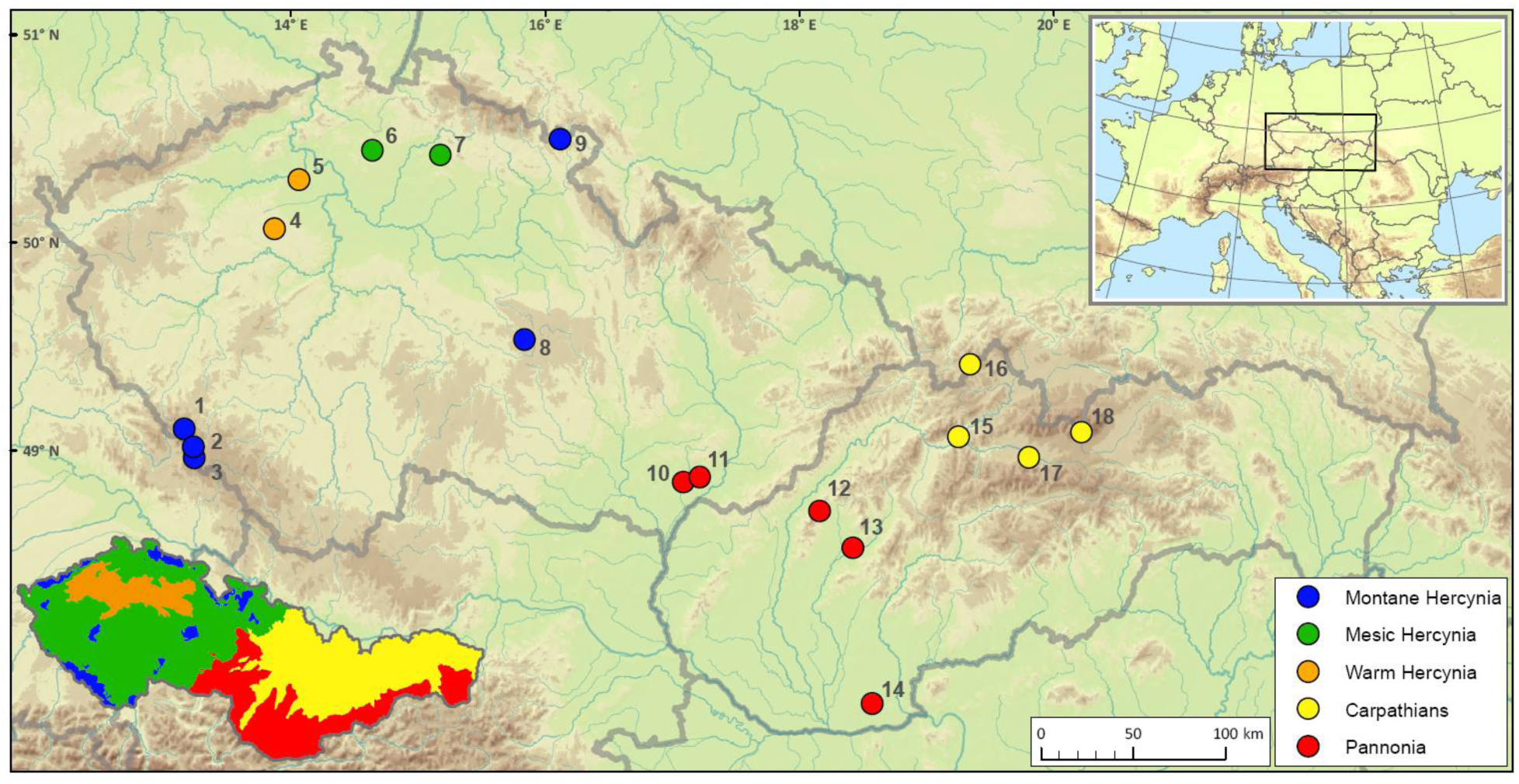
Study area with study sites and their classification to biogeographical regions shown. Study sites: 1 – Hůrecká slať; 2 – Stará Jímka; 3 – Rokytská slať; 4 – Rynholec; 5 – Zahájí; 6 – Okna; 7 – Čin-Čan-Tau; 8 – Dářko; 9 – Velké ohbí; 10 – Svatobořice-Mistřín; 11 – Vracov; 12 – Mitická slatina; 13 – Bielice; 14 – Paríž; 15 – Stankovany; 16 – Zlatnická dolina; 17 – Liptovský Ján; 18 – Popradské pleso. See Table 1 and Appendix S1 for details on study sites.

Eastern part of the study area is formed by Western Carpathians Mts. in the north and Pannonian lowland in the south. The Carpathians are an arc-shaped mountain belt formed in the Late Mesozoic and the Tertiary. Their younger origin results in high topographic complexity. Calcareous bedrock is common also in higher altitudes. Large intermontane basins are a characteristic feature of the region. Southerly, lower mountain ranges with diverse bedrock border the Pannonian Basin. While the mountains generally have a moist and cold climate and remain largely forested up to the treeline, the intermontane basins have a rather continental climate and a long history of open landscape connected with agricultural use (Soják, 2001; Soják & Furman, 2018).

**Table 1.** Basic information on geography and environmental conditions of the study sites. Additional information is available in Appendix S1.

| Site name | latitude<br>[°N] | longitude<br>[°E] | elevation<br>[m a.s.l.] | mean<br>annual<br>temperature<br>[°C] | mean<br>annual<br>precipitation<br>[mm] | Biogeographical<br>region |
| --- | --- | --- | --- | --- | --- | --- |
| Bielice | 48.629 | 18.34378 | 190 | 10.0 | 581 | Pannonia |
| Čin-Čan-Tau | 50.51811 | 15.20271 | 265 | 8.4 | 683 | Mesic Hercynia |
| Dářko | 49.63787 | 15.87563 | 635 | 7.1 | 667 | Montane Hercynia |
| Hůrecká slať | 49.15222 | 13.32755 | 870 | 6.0 | 903 | Montane Hercynia |
| Liptovský Ján | 49.04167 | 19.67778 | 660 | 6.7 | 854 | Carpathians |
| Mitická slatina | 48.80833 | 18.10028 | 318 | 9.0 | 649 | Pannonia |
| Okna | 50.53207 | 14.67593 | 280 | 8.6 | 586 | Mesic Hercynia |
| Paríž | 47.87494 | 18.46144 | 123 | 10.6 | 508 | Pannonia |
| Popradské pleso | 49.1535 | 20.07984 | 1494 | 1.4 | 1079 | Carpathians |
| Rokytská slať | 49.0153 | 13.4122 | 1097 | 4.7 | 1332 | Montane Hercynia |
| Rynholec | 50.13719 | 13.94188 | 410 | 8.0 | 478 | Hercynian Warm |
| Stankovany | 49.1534 | 19.1534 | 437 | 7.8 | 790 | Carpathians |
| Stará Jímka | 49.06889 | 13.40278 | 1110 | 4.6 | 1232 | Montane Hercynia |
| Svatobořice-Mistřín | 48.95432 | 17.08188 | 174 | 10.1 | 472 | Pannonia |
| Velké ohbí | 50.6045 | 16.12841 | 583 | 6.6 | 717 | Montane Hercynia |
| Vracov | 48.97784 | 17.2052 | 193 | 10.0 | 516 | Pannonia |
| Zahájí | 50.37922 | 14.11555 | 232 | 8.9 | 503 | Warm Hercynia |
| Zlatnická dolina | 49.5 | 19.25833 | 850 | 5.3 | 987 | Carpathians |

Pannonian (Carpathian) Basin is a large basin enclosed between the Alps, the Carpathians and the Dinarides (Rasser & Harzhauser, 2008). It originated during the Alpine Orogeny and is filled with mostly calcareous sediments of Tertiary age, usually overlain by fluvial sediments, loess or eolian sands. The climate of the basin is relatively warm and continental, with rather low precipitation sums with maxima during the summer. There is a rich history of human activities and extensive agricultural use since the Neolithic (Tóth et al., 2011).

### Data acquisition

The study area is densely covered with pollen profiles (Kuneš et al. 2009; https://botany.natur.cuni.cz/palycz). However, the records available in the databases are inconsistent in terms of data quality. Therefore, a strict selection was applied prior to the analysis: only sites covering long periods of the Holocene (at least 9000 years, often reaching the Late Glacial), analysed with the highest effort, precision and taxonomic resolution, and with a reliable depth-age model available, were involved. Altogether 18 high-quality fossil pollen profiles (Table 1 and Appendix S1) were selected; one of them was newly analysed for this purpose (Roleček et al., 2020).

### Data analysis

We used two main descriptors of the Holocene diversity pattern: mean pollen richness per site and variation in pollen richness on different time scales. To account for biases in pollen richness, pollen assemblages with harmonized taxonomy were standardized for equal detection probability and sampling effort. We had new pollen productivity estimates for one of the study regions available (White Carpathian forest steppe; Kuneš et al. 2019), however their applicability outside the region is uncertain. Therefore, we used averaged pollen productivity values for the Northern Hemisphere (Wieczorek & Herzschuh, 2020), which should provide robust estimates suitable for extrapolation to the past. We divided pollen counts of tress by corresponding pollen productivity values (see Appendix S2). Subsequently, adjusted pollen counts were resampled to 100 grains per sample and 150 grains per 300-years time window. Each time window was resampled 100 times, and pollen richness was recorded. Sample or time window with median richness was used for further analyses. Mean site richness during the Holocene was calculated as the mean of the time-window richness values available for each profile.

SiZer analysis was used to identify significant patterns in Holocene diversity variation. SiZer is a multi-scale polynomial regression analysis producing graphical output called SiZer map (Chaudhuri & Marron, 1999; Park et al., 2012). It is a two-dimensional image that summarises the positions of all the statistically significant slopes, where these slopes are estimated by smoothing the data with different resolution (bandwidth). Time windows on X-axis were set to 300 years. Bandwidths on Y-axis ranged from 500 to 5000 years with 41 logarithmic increments in between. We evaluated the similarity of SiZer maps using non-metric multidimensional scaling (NMDS) ordination. For this purpose, we calculated an agreement matrix for each pair of SiZer maps. We scored the match of the 1681 corresponding cells of SiZer maps as follows: both cells showed the same trend, positive or negative: 2 points; both cells showed no trend: 1 point; lack of data in one or both cells: 0 points; one cell showed no trend, the other positive or negative trend: -1 point; corresponding cells showed opposite trends: -2 points. In the end, we summed these values for each pair of maps and transformed the resulting matrix to a dissimilarity matrix by dividing the values by the theoretical maximum (i.e. 3362) and subtracting the quotient from 1. We reduced the information in the dissimilarity matrix to a single dimension using NMDS in PC-ORD 6.0 (McCune & Mefford, 2011), with 500 iterations and number of axes set to 1. The resulting values for each site may be simply compared and analysed, and we interpret them as positions on the main gradient in diversity dynamics.

To explain the identified patterns in mean diversity and temporal diversity dynamics, we assessed which geographical and environmental factors predict best their variation using a linear model in R (R Core Team, 2019). We tested latitude, longitude, elevation, mean yearly temperature and mean yearly precipitation. Because our study area is biogeographically complex, we also included classification of the study sites to main biogeographical regions. All predictors except biogeographical regions were scaled and centred before the analysis. Based on the Akaike information criterion, a stepwise algorithm was used to select the final model. Temperature and precipitation data were retrieved from CHELSA dataset (Karger et al., 2017). Delimitation of biogeographical regions was adopted from the phytogeographical division of the Czech Republic (Skalický, 1988) and Slovakia (Futák, 1966). Five regions were defined as follows (see Figure 1 and Kaplan 2012): Warm Hercynia (Bohemian Thermophyticum), Mesic Hercynia (Bohemian-Moravian Mesophyticum), Montane Hercynia (Bohemian-Moravian Oreophyticum), Carpathians (Carpathian Oreophyticum in the Czech Republic and Carpaticum occidentale in Slovakia), and Pannonia (Pannonian Thermophyticum in the Czech Republic and Pannonicum in Slovakia).

To relate the identified diversity patterns to compositional changes in fossil pollen assemblages during the Holocene, we performed NMDS of fossil pollen compositional data. We projected a smooth surface of corresponding pollen richness values over an ordination plot using *ordisurf* function from *vegan* library in R (Oksanen et al., 2019). Abundances were transformed to presence-absence scale to reflect as much as possible of the richness changes. To show the geographical pattern, we grouped the sites by biogeographical regions. Thanks to this grouping, we were able to include also the Late Glacial period, for which data were available only for some sites. To assist the interpretation of ordination results, we projected centroids of samples grouped by K-means clustering (presence-absence abundance scale, 100 cycles, seven clusters) onto the ordination plot in R. Composition of the clusters was characterised using the lists of dominant, constant and indicator pollen taxa. Indicator value of taxa was measured with the phi coefficient of fidelity standardised to group size equal to 15% of the total dataset (Tichý & Chytrý, 2006).

To generalise our results, we calculated mean diversity curves for the sites grouped by biogeographical regions. We calculated mean pollen richness for each time window and each biogeographical site group.

### Testing the pollen-plant diversity relationship

At two study sites (Dářko and Vracov), we tested the assumption that fossil pollen is informative in terms of past plant diversity in our study area. First, we compared pollen richness in the last 1000 years with modern pollen richness in the reference areas: the Bohemian-Moravian Highlands (a species-poor region with coniferous forests and wet meadows) for Dářko site and the White Carpathians (a species-rich region with broad-leaved forests and steppe meadows) for Vracov site. Second, we compared numbers of modern pollen and plant taxa in the reference areas using extensive field sampling. Data for the distance where the overall regression between recent pollen and plant richness was highest was used: 550 m in the Bohemian-Moravian Highlands (R^2^ = 0.52) and 250 m in the White Carpathians (R^2^ = 0.51). The interaction with habitat (forested versus open) was tested using ANOVA. The methodology of sampling of modern pollen and floristic data and specific analyses will be published in a separate study (Abraham et al., 2020).

## Results

### Patterns in Holocene plant diversity

Reconstructed mean Holocene plant diversity and position on the main gradient in diversity dynamics for all study sites are shown in Table 2 and Figure 2. Sites in Mesic and Montane Hercynia are clearly separated from the sites in Pannonia, the Carpathians and Warm Hercynia. The linear model analysis showed that latitude is the only significant predictor of variation in mean diversity between the sites (ANOVA, F-statistic: 7.97, p-value: 0.012). The position on a single NMDS axis, corresponding to the main gradient in diversity dynamics, was significantly influenced by biogeographical region (ANOVA, F-statistic: 11.51, p-value < 0.01) and elevation (ANOVA, F-statistic: 7.79, p-value: 0.02). Significant differences were found between Pannonia, the Carpathians and Warm Hercynia on the one hand and Mesic and Montane Hercynia on the other hand (Appendix S3). Diversity curves and SiZer maps for all sites are provided in Figure 3. They show that the main gradient in diversity dynamics separates sites with diversity increasing only in the Late Holocene (diversity trend is then often close to unimodal) and sites with diversity increasing already in the Early or Middle Holocene (diversity trend is then often close to linear positive or more complex). NMDS ordination of compositional data (Figure 4) and analysis of indicator species (Appendix S4) show that low diversity in the Middle Holocene and at the beginning of the Late Holocene, characteristic for the Mesic and Montane Hercynia, is associated with the prevalence of competitive woody taxa (mainly *Tilia, Ulmus, Fraxinus, Quercus, Corylus* and *Picea* in the Middle Holocene and *Fagus, Abies* and *Carpinus* in the Late Holocene) and acidophilic herb and low shrub taxa (e.g. Vacciniaceae, *Calluna, Melampyrum*). Higher diversity in Pannonia, the Carpathians and Warm Hercynia is associated with the presence of many herb taxa, including grassland and minerotrophic wetland taxa, crops and weeds (e.g. *Alisma, Centaurea*, Cerealia, Chenopodiaceae, Compositae, *Daucus, Nymphaea, Plantago major*/*media, Polygonum aviculare, Sparganium*), while woody taxa composition may be similar to the previous group of sites. The identified trends in plant diversity are summarised using locally weighted regression curves for sites grouped by biogeographical regions (Figures 5 and Appendix S5).

**Table 2.** Basic characteristics of the Holocene diversity pattern for all study sites. The sites are sorted by increasing diversity. Position on the main gradient in diversity dynamics is a position on a single axis of NMDS ordination of SiZer maps (see Figure 3).

| Site name | Mean diversity | Position on the main gradient in diversity dynamics |
| --- | --- | --- |
| Hůrecká slať | 14.07 | -1.13035 |
| Čin-Čan-Tau | 14.16 | -0.83801 |
| Dářko | 15.02 | -1.75905 |
| Velké ohbí | 15.43 | -0.02243 |
| Rokytská slať | 15.56 | -1.52873 |
| Rynholec | 18.11 | 1.16112 |
| Liptovský Ján | 18.98 | 1.52526 |
| Zahájí | 19.10 | 0.61469 |
| Svatobořice-Mistřín | 19.40 | 0.68495 |
| Stará jímka | 19.69 | -0.97104 |
| Stankovany | 19.72 | 0.30408 |
| Zlatnická dolina | 19.98 | 0.12105 |
| Okna | 20.38 | -1.55323 |
| Mitická slatina | 20.55 | 0.85251 |
| Popradské pleso | 21.30 | 0.43136 |
| Vracov | 21.48 | 0.33424 |
| Bielice | 24.75 | 0.94478 |
| Paríž | 24.82 | 0.82880 |

**Figure 2.**
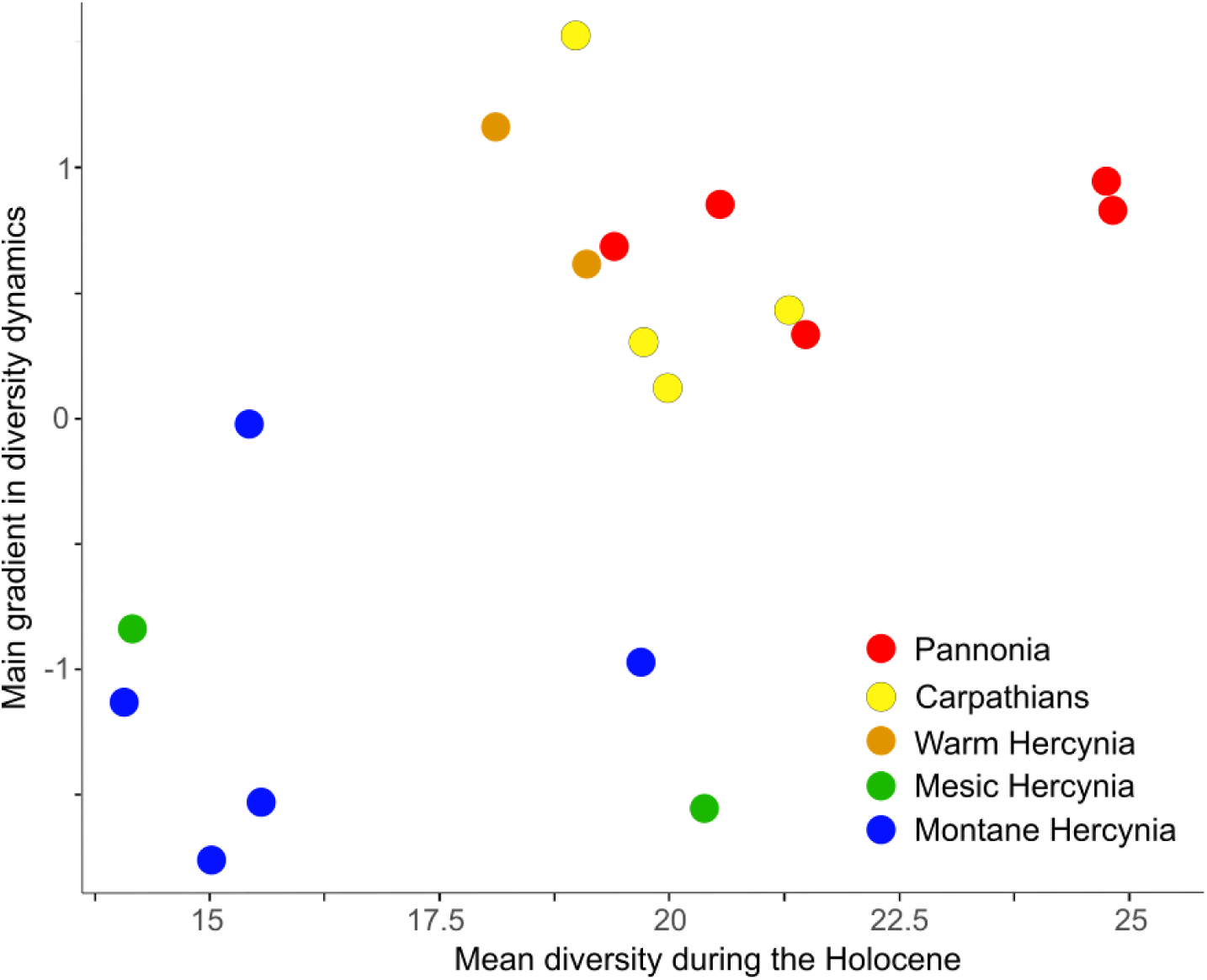
Relationship between mean diversity and position on the main gradient in diversity dynamics. Colours of symbols indicate different biogeographical regions.

**Figure 3.**
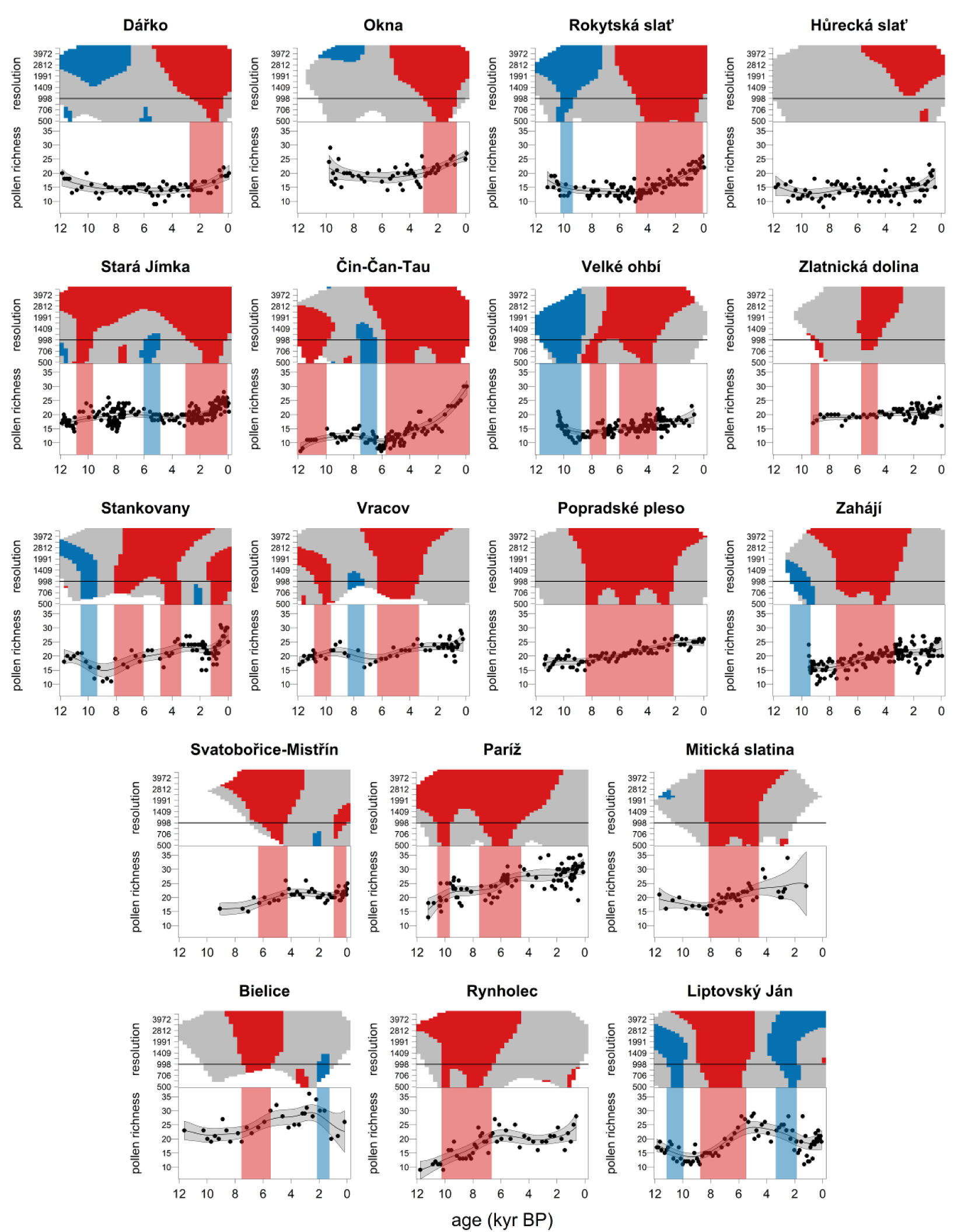
SiZer maps (above) and pollen richness curves (bellow) for all analysed sites. Red and blue colours indicate increasing and decreasing diversity, respectively; grey in SiZer maps indicates no trend, white insufficient data. The Y-axis of SiZer maps shows temporal resolution (bandwidth) for which the trend was calculated. Locally weighted polynomial regression at bandwidth 998 years was used to draw the pollen richness curves. The graphs are arranged from top left to bottom right according to the position on a single NMDS axis (Table 2). The gradient separates sites where diversity increases only in the Late Holocene from sites where diversity increases already in the Early or Middle Holocene.

**Figure 4.**
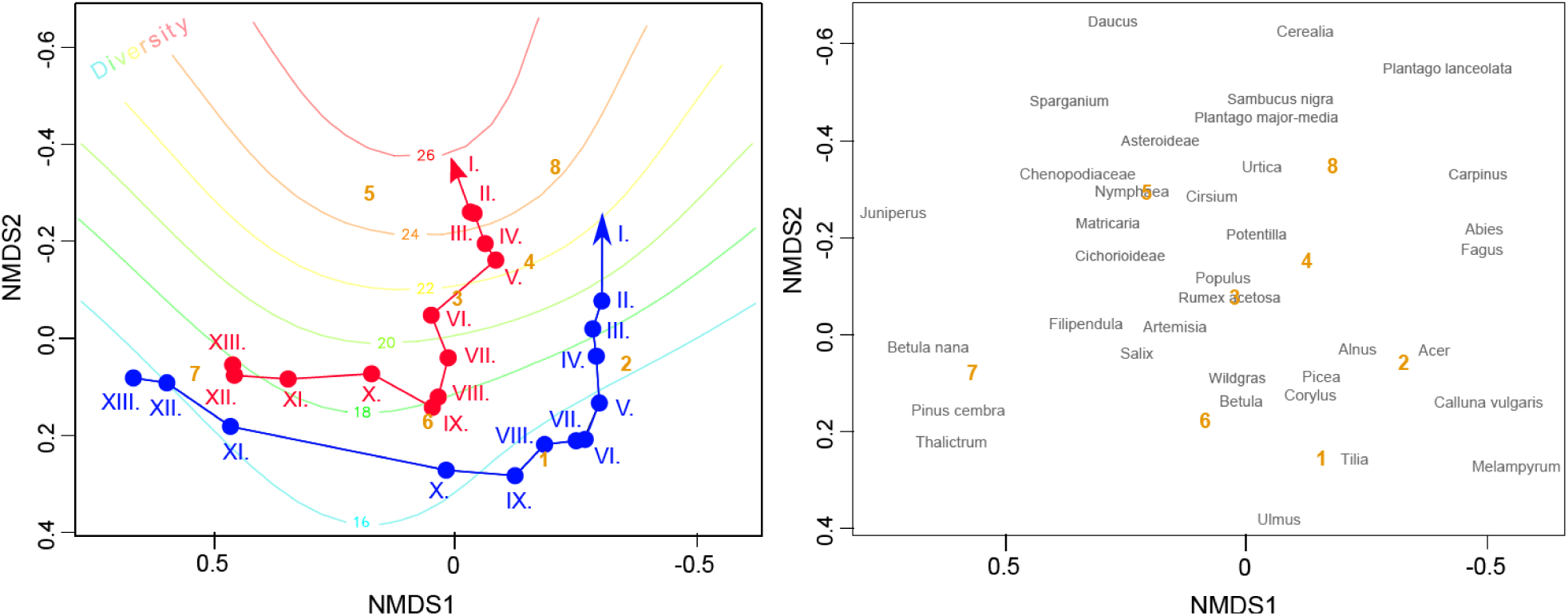
NMDS plot showing the relationship between plant diversity and changes in species composition during the Holocene. Left: Arrows connect centroids of samples of different age (nodes and Roman numerals indicate millennia BP). Colours of arrows indicate different biogeographical regions (red: Pannonia, the Carpathians and Warm Hercynia; blue: Mesic and Montane Hercynia). Corresponding diversity values were fitted with a generalised additive model and projected on the ordination plot as isolines. Bold orange numbers 1–8 are centroids of samples grouped by K-means clustering, whose indicator taxa are provided in Appendix S4. Right: Taxa with frequency > 5% are shown; if several labels overlap, the more frequent taxon is plotted.

**Figure 5.**
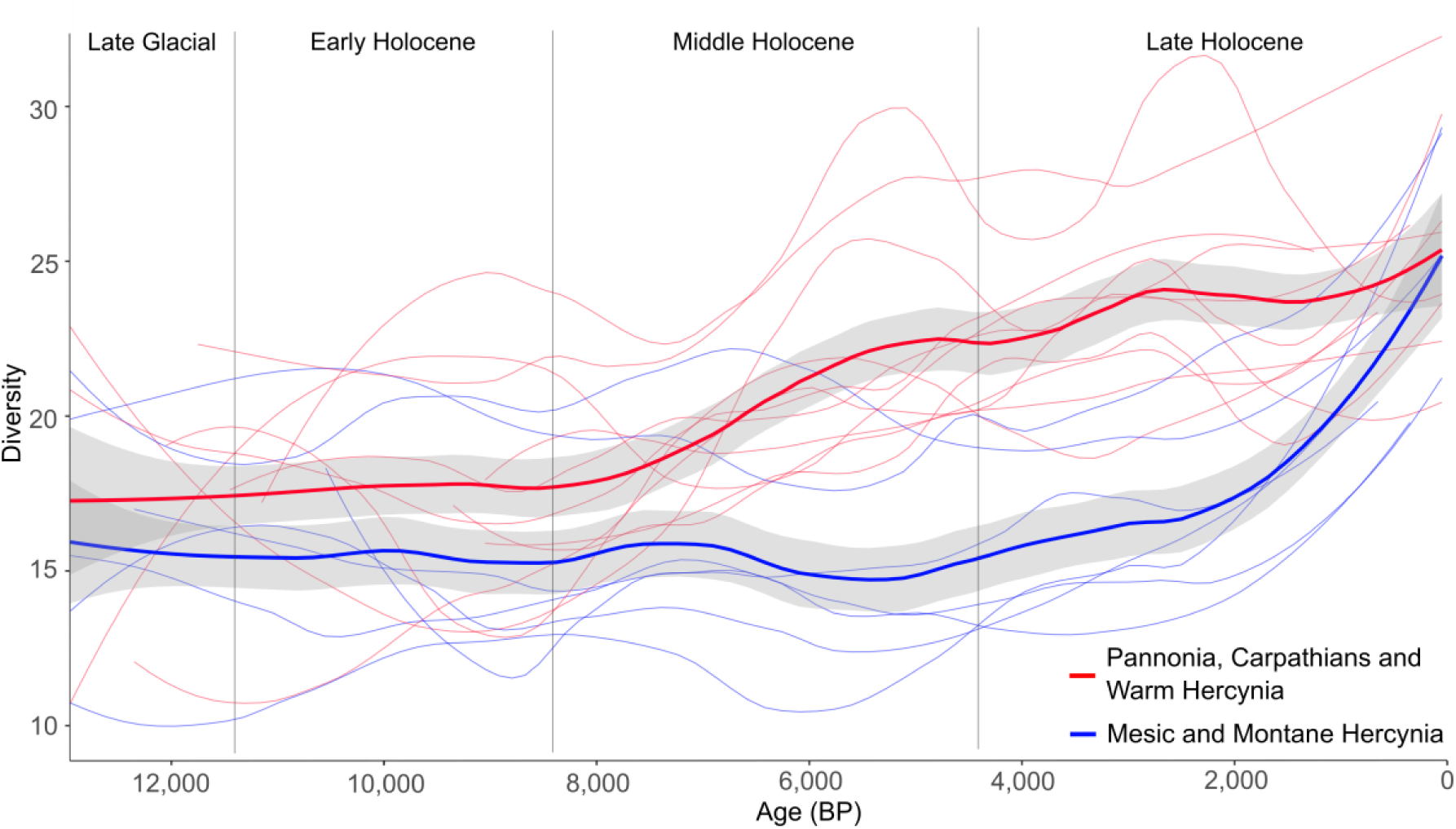
Two main Holocene trends in pollen-based plant diversity in the study area. Locally weighted polynomial regression fitted to pollen richness values grouped by biogeographical regions is shown. Grey band indicates 95% confidence interval. Level of smoothing was set to 0.4. Thin lines are regression curves for individual study sites. Differences between the biogeographical regions are statistically supported (see the linear model test in the Results and Appendix S3).

### Fossil pollen as a proxy of plant diversity

There was no significant difference in diversity between sub-recent fossil pollen (last 1,000 years) of Vracov site and modern pollen of its reference area, the White Carpathians (means 23.1 and 23.9; ANOVA, F-statistic: 0.598, p-value: 0.443) or recent fossil pollen of Dářko site and modern pollen of its reference area, the Bohemian-Moravian Highlands (means 19.2 and 18.5; F-statistic: 0.306, p-value: 0.585). At the same time, there was a significant difference in diversity of recent fossil pollen between Dářko and Vracov sites (ANOVA, F-statistic: 6.857, p-value: 0.018). See also Appendix S6.

We found a significant relationship between recent plant and pollen diversity in both the White Carpathians (ANOVA, F-statistic: 41.63, p-value < 0.01) and the Bohemian-Moravian Highlands (ANOVA, F-statistic: 22.57, p-value < 0.01), with no difference between the open and forested habitats in both regions (ANOVA, F-statistic: 0.34, p-value: 0.56 for the White Carpathians and F-statistic: 0.40, p-value: 0.53 for the Bohemian-Moravian Highlands).

## Discussion

### Holocene plant diversity patterns mirror latitude, elevation and biogeographical regions

Distribution of biological diversity on the Earth shows clear patterns, sometimes even considered rules. Among them, the latitudinal diversity gradient (LDG) is one of the most robust (Hillebrand, 2004; Mittelbach et al., 2007). The mechanisms behind LDG are probably complex (Wiens & Donoghue, 2004), and it is affected by factors such as longitude and habitat (Kinlock et al., 2018). Giesecke et al. (2019) have shown that LDG is visible also in the fossil pollen record and described its Holocene dynamics across Europe. They suggested that the gradient has strengthened during the Holocene due to the combined effects of climate and human activities, with the latter factor being more important. šizling et al. (2016) even refused the influence of temperatures on the Holocene diversity trends in Central Europe and emphasised the human impact.

Our results support the view that mechanisms behind the temporal diversity pattern within a single region are complex. Although we also identified latitude as the best predictor of the mean Holocene plant diversity, we found a finer-scale structure when analysing Holocene diversity dynamics. The identified patterns can be explained in terms of geographical position, biogeographical relationships, and past changes in plant assemblages and landscape openness. They differentiate the middle and higher elevations in the north-western part of the study area (Hercynia) from the warm and dry lowlands plus the Carpathian mountain range. Diversity was low during most of the postglacial in the former region and increased sharply only during the Late Holocene (typically since the Medieval Age). On the contrary, diversity was higher in the latter region and increased gradually from the beginning of the Holocene, with a marked acceleration in the Middle Holocene.

Our results largely conform to the classic concept of two developmental pathways, proposed for Central-European landscapes based on the study of Holocene succession of mollusc assemblages (Horsák et al., 2019; Ložek, 1982). The main difference we identified in plant diversity dynamics is that the Carpathians are more similar to lowlands across the study area than to the Hercynian mountain ranges (see possible explanations below).

We also want to acknowledge the authors of the phytogeographical division of the study area (Futák, 1966; Skalický, 1988). We believe our results support their concepts and reinforce the view that the present diversity of Central European flora reflects its diverse Holocene histories.

### Drivers of the geographical patterns

The fact that biogeographical regions significantly structure the observed patterns allows seeking explanations for the patterns in the known differences between the regions. As biodiversity of an area depends on multiple drivers including quality and quantity of available resources, species interactions and historical factors such as speciation, extinction and migration (Diamond, 1988; Whittaker & Fernández-Palacios, 2007), relevant differences between the regions may be found e.g. in habitat conditions (including climate), sequence of Holocene compositional changes of biotic assemblages, refugial history, or history of human influence. In the Hercynian mountain ranges, the postglacial vegetation developed in general from species-poor to moderately rich mosaic of taiga, tundra, and steppe in the Late Glacial, towards very species-poor, densely-wooded landscape dominated by temperate mixed forests during the Holocene. *Picea* played an important role here (Kuneš & Abraham, 2017). Relatively high precipitation and low temperatures supported the development of closed forests, which peaked by the end of the Middle Holocene, when highly competitive broadleaved trees (*Fagus, Abies, Carpinus*) took over the dominance followed by the Holocene diversity minimum. In the Late Holocene, diversity started to increase, apparently in connection with deforestation and landscape transformation by humans, as indicated by an increase in human indicators (grassland taxa, crops, weeds). Low diversity in the Hercynian mountain ranges was probably also supported by lower topographic complexity compared to the Carpathians, and prevalence of acidic soils with restricted evolutionary species pool (Ewald, 2003). Acidity might have been enhanced by leaching connected with high precipitation during the Middle Holocene, and in some places also by a high representation of conifers, either induced by human activities or of natural origin (Juřičková et al., 2020; Novák et al., 2012). Hercynian lowlands, having warmer and drier climate and mostly base-rich soils, remained relatively more open throughout the postglacial, with higher proportions of oak and pine (Abraham et al., 2016; Pokorný et al., 2015). Their diversity dynamics is more similar to that of the Pannonian lowlands.

Pannonian region became relatively species-rich already during the Early Holocene, probably due to drier and warmer climate and prevalence of base-rich soils that allowed continuous persistence of open habitats (forest steppe) from the Last Glacial period (Kuneš et al. 2015). With climatic amelioration at the beginning of the Holocene, temperate and sub-Mediterranean species have spread, and this process might have been accelerated by the proximity to presumed glacial refugia of temperate species (Jamrichová et al., 2014). The adjacent Carpathian region, however, experienced an early spread of forests with the dominance of canopy trees with high shading ability (*Picea, Tilia, Ulmus*; Abraham et al., 2016; Hájek et al., 2016), probably due to moister climate. The simultaneous marked decrease in diversity by the end of the Early Holocene (see Appendix S5) may thus result from the fast-closing canopies (see also Hájek et al., 2016). In the Middle Holocene, both lowland regions and the Carpathians show a remarkable increase in diversity. In agreement with previous studies (Giesecke et al., 2019; šizling et al., 2016), we see the main reason in the early development of diverse agricultural landscape, conditioned by the immigration of Neolithic farmers around 7,500 BP (Bramanti et al., 2009). While this explanation is evident for the Pannonian lowlands with deep, base-rich, fertile soils suitable for agriculture (Tóth et al., 2011), it may also apply to the Inner Carpathian basins (Soják, 2001; Soják & Furman, 2018). Sources of the increasing plant diversity were varied, as indicated by a rich spectrum of indicator taxa, including grassland and minerotrophic wetland taxa, crops and weeds.

### Fossil pollen as a proxy of plant diversity

We examined the predictive value of pollen as a proxy of past plant diversity and showed that it may be useful if treated appropriately. Comparisons of both fossil-modern pollen diversity and modern pollen-plant diversity suggest that reduction of tree influence on pollen sum is a key analytical step. Interestingly, pollen richness in modern pollen assemblages adjusted to tree pollen productivity did not show better correlation with floristic richness (Abraham et al., 2020), probably because studied landscapes are presently dominated by low pollen producers (Kuneš et al., 2019). On the other hand, present boreal forests (Felde et al., 2016) and Early Holocene forests in our study area host(ed) several high pollen producers (*Pinus, Betula*) decreasing detection probability of rare pollen taxa. After removing this bias by using up to date pollen productivity estimates for trees (Wieczorek & Herzschuh, 2020), the adjusted modern pollen assemblages have not shown significant difference in diversity between forested and open sites any more. This result indicates the reduction of the effect of local high pollen producers and the manifestation of a shared regional pollen pool.

We think that our methodological framework may be useful in further research of past plant diversity. At the same time, it requires further testing, particularly concerning its generality, as the relationships between plant and pollen diversities may differ in different ecosystems (Birks, Felde, Bjune et al., 2016). Such differences may have consequences even for our study area, particularly for the reconstruction of diversity changes in more distant past (LGM, Late Glacial), when plant and pollen assemblages were composed differently (Reille et al., 2000). Calibration studies from a broad array of ecosystems are therefore needed.

## Acknowledgements

Our thanks go to Barbora Werchan for help with sampling recent pollen data, Pavel Daněk, Helena Prokešová, Kryštof Chytrý and Zita Červenková for help with sampling plant diversity data, Ondřej Hájek for GIS support, Stano Pekár, Petr šmilauer and David Zelený for advice on statistics, Michal Hájek and Petra Hájková for information on some Carpathian sites, and the staff of Tisůvka pension for providing a pleasant field base. This study was supported by the Czech Science Foundation (Grant No. 16-10100S). Authors affiliated to the Institute of Botany were further supported by the long-term developmental project of the Czech Academy of Sciences (RVO 67985939).

## Author contributions

J.R., V.A. and P.K. conceived the ideas; J.R., O.V., Z.P., V.A., E.J., H.S.S. and P.P. collected the data; V.A., J.R. and O.V. analysed the data; and J.R. led the writing. All authors commented upon the manuscript.

